# Directionality range in Emlen funnels

**DOI:** 10.1101/2025.02.04.636393

**Authors:** Ilias Patmanidis, Bo Leberecht, Martin Fränzle, David Lentink, Ilia A. Solov’yov, Henrik Mouritsen

## Abstract

Emlen funnels can be used to study the birds’ ability to orient during the migratory seasons. Birds are so eager to migrate that they will jump in the direction in which they want to fly, even if they are placed in small cages during the night. Emlen funnels have therefore been used for decades to study the sensory capabilities and mechanisms that migratory birds use to find their way. A significant part of this research has focused on how night-migratory songbirds perceive the Earth’s magnetic field. Even though Emlen funnels have been proven quite successful in capturing the birds’ behavioural responses to different experimental conditions, the orientation behaviour of night-migratory songbirds in Emlen funnels is very noisy, i.e. tends to have a low signal to noise ratio. This noise makes Emlen funnel experiments very time consuming and limits the types of questions that can be studied to those which require very few different experimental conditions (permutations). Furthermore, the experimental design choices can be crucial, e.g. degree of blinding of the experimenters and data evaluators to the conditions tested, pre-testing of birds to make sure they are in a migratory state, and planning of the effective sample sizes. Hence, different traditions in experimental design choices can reduce reproducibility and comparability and render the interpretation of the results non-trivial. To better understand Emlen funnel data and the minimal requirements for good experimental design, we constructed and analyzed a large data set that we compiled by combining behavioural data from many previous Emlen funnel studies performed in Oldenburg. Our results provide realistic ranges for the expected orientation of the birds in Emlen funnels, which can be useful (and in some research crucial) for predicting the optimal sample sizes for future experiments. Our results thus offer concrete information for the design and analysis of statistically powerful future magnetic orientation experiments.

## Introduction

Bird migration and the mechanisms that birds use to orient have attracted the interest of scientists for many decades. ^1?–9^ Years of research have shown that birds can use many different cues to guide their migratory journeys, such as geographical landmarks, celestial visual information (e.g. the sun and the stars), the Earth’s magnetic field, and even social interactions.^1? –11^ Temperature and weather can also influence bird migration, as they, for instance, influence the motivation of birds to take off and migrate. ^12–15,15–19^

Magnetoreception, the ability to detect and respond to the geo-magnetic field, plays a crucial role in bird migration and avian navigation.^6,9,20–23^ While the exact physico-chemical mechanisms of magnetoreception are not fully understood, its use is demonstrated by ample behavioural evidence. ^3,9,24^ The leading hypothesis assumes the photo-activation of light-sensitive cryptochrome 4 proteins in the avian retina and the consequent formation of radical pairs that are sensitive to the Earth’s magnetic field.^23,25–29^ The magnetic field sensitivity of the radical pairs is expected to be transmitted to the brain via the thalamofugal visual pathway.^20,30–34^ The radical pair formation causes a cascade of events starting with changes in the cryprochrome’s electron spin dynamics which ultimately affects the bird’s orientation behaviour. ^28^ Alternative hypotheses suggest either magnetic particles (e.g. magnetite or maghemite) acting like microscopic compass needles^35,36^ or sensation of magnetic induction in the semicircular canals. ^37,38^ The changes in the orientation behaviour based on either of these hypothesised magnetic perception mechanisms has been studied in Emlen funnels. ^39^

Behavioural experiments in Emlen funnels^39–42^ are a key paradigm for studying bird orientation behaviour and understanding the magnetic sensory mechanism(s) of birds. In these experiments, a bird is placed into a funnel-shaped cage lined with scratch-sensitive paper on its walls (an improvement upon the first inkpad-and-paper-based setup).^41^ Since the birds, on average, tend to jump towards their preferred migratory direction, their intended orientational direction and their ability to sense magnetic fields is evaluated by analysing the scratches left by the birds’ feet on the scratch-sensitive paper, Figure 1A. By blocking visual orientation cues, birds can be tested while they only rely on their magnetoreception capabilities to determine their preferred direction. ^32,43?–45^ Furthermore, Emlen funnels provide a controlled environment, in which potential disturbances to their sensory systems can be tested such as the effects of radio frequency electromagnetic noise (also called electro-smog) on the birds’ magnetic compass orientation capabilities in the context of the radical pair mechanism of avian magnetoreception. ^44–51^

**Figure 1:**
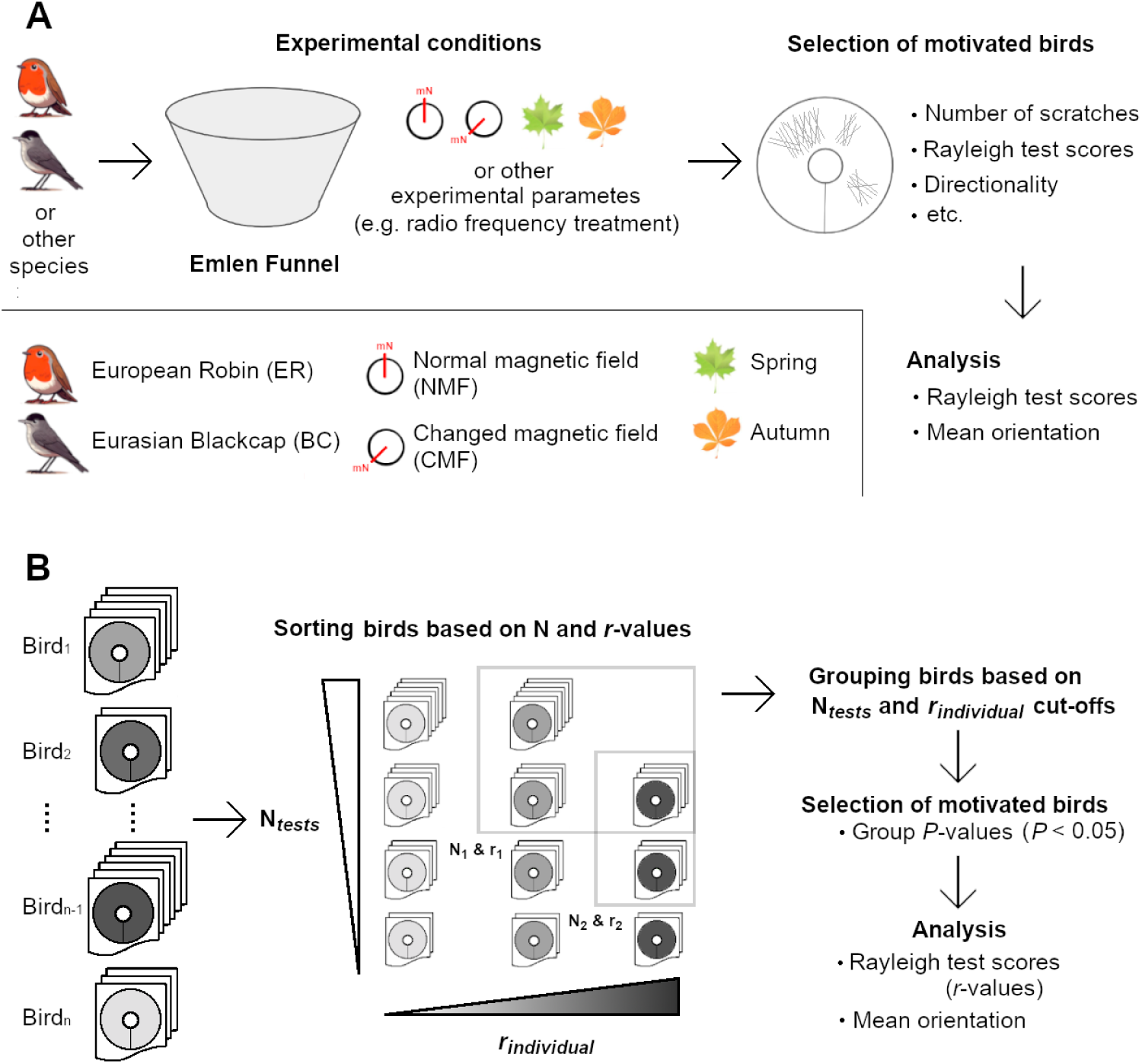
A: Schematic representation of the workflow in Emlen funnel experiments. B: Schematic representation of our data analysis workflow of the compiled dataset used in this work. The birds are categorized based on the number of experiments/scratched papers (N*_tests_*, number of tests per individual) that they participated in and their associated individual directedness (*r_individual_* from the Rayleigh test). All birds with larger values than the set N*_tests_* and *r_individual_* cut-offs were grouped together and the deviation from uniformity was determined using the *P* -value from the Rayleigh test. For each such group, its directedness (*r*) and mean orientation were calculated. Effectively the groups with high cut-offs include the most consistently oriented birds and the birds with more tests.

While Emlen funnels have provided a lot of very important knowledge about the sensory systems and mechanisms birds used for navigation, they also have some methodological weaknesses. In nature, birds do not migrate every day: they mix stopover time for resting, recovering and refuelling with migration days/nights. ^18^ One important consequence of this is that birds are not motivated to migrate every time that they are put in an Emlen funnel. Consequently, on some test nights, the birds tested in Emlen funnels are expected to be oriented in random directions (non-motivated bird nights), while on other nights, they are expected to be oriented mainly in the mean migratory direction (motivated bird nights). The challenge is that there is no way to determine *a priori* whether a given bird is motivated to migrate on a given night. Consequently, the behavioural recordings in Emlen funnels tend to contain noisy data whose interpretation is not straightforward, especially when the sample sizes are too low. The way most researchers using Emlen funnels deal with this noise is to test the birds enough times, so that the motivated, oriented nights are likely to be reflected in the mean direction of many tests of a given bird. Hence, different protocols for conducting Emlen funnel experiments and analyzing the behavioral data have been proposed. ^32,40–44,52^**^?^** ^,53^

The present exploratory study combines data from a large collection of Emlen funnel experiments and aims to quantify general trends that govern the consistency of bird orientation in the Emlen paradigm. We combined data sets from magnetic-cues-only studies performed by researchers from the University of Oldenburg using very similar protocols for all studies. Our results provide realistic ranges for the expected magnetic orientation of the birds in Emlen funnels and highlight the statistical limitations of commonly used analyses.

Critically, our results offer data-informed experimental insights for designing and analyzing future magnetic orientation experiments in Emlen funnels. Apart from the fundamental questions regarding avian navigation and migration, understanding and parametrizing the underlying orientation mechanisms is also important for the development of biomimetic navigation systems that mimic the sensory abilities of birds. ^54–56^ Ultimately, similar navigation principles can be utilized to design novel technologies and algorithms for the navigation and orientation of artificial robotic devices.

## Methods

### Data set

We analysed data from Emlen funnel studies dating from 2009 to 2023. ^45,47,49–51,57–60^ We gathered the raw digital data of all the studies from 2009 to 2023 and merged them to all conform to the same formatting. ^45,47,49–51,57–60^ The entries were spell checked and verified by comparing them to the lab books and spreadsheets of the respective migratory seasons to ensure that the analysis of each scratch paper was assigned correctly (for 194 entries parts of the scratch paper analysis could not be retrieved). While all birds had numbered colour rings, occasionally combinations of number and colour were re-used or changed. To track the reoccurring and changing number-colour-combinations, all individual birds were distinctly labelled by comparing their numbered colour rings to arrival and departure dates in the animal keeping and catching logs. To get the most realistic picture of the noise in Emlen funnel data, our dataset includes data that were collected during the conduct of these studies, but in which the control conditions did not result in significant group orientation. In these cases, the data could not be clearly interpreted and, therefore, remained unpublished. Lastly, the data was filtered to only contain orientational values for control conditions of European robins and Eurasian blackcaps, i.e. removing entries evaluated as inactive, randomly oriented, or bimodally oriented.

In total, we collected 27.448 curated entries for the preferred bird orientation based on scratched papers from Emlen funnels, Figure 2. This is the largest laboratory-based data set that has so far been compiled and analysed in detail to study the magnetic orientation of migratory birds and the statistics of Emlen funnel data. The studies were performed on two migratory species, European robins and Eurasian blackcaps, for two different magnetic field conditions, normal magnetic field (NMF) and changed magnetic field (CMF). In the CMF condition, the horizontal component of the magnetic field was rotated 120*^◦^* counter-clockwise. In addition to species and magnetic conditions, the data set included information on the bird activity, the concentration of scratches and the date of each measurement.

**Figure 2:**
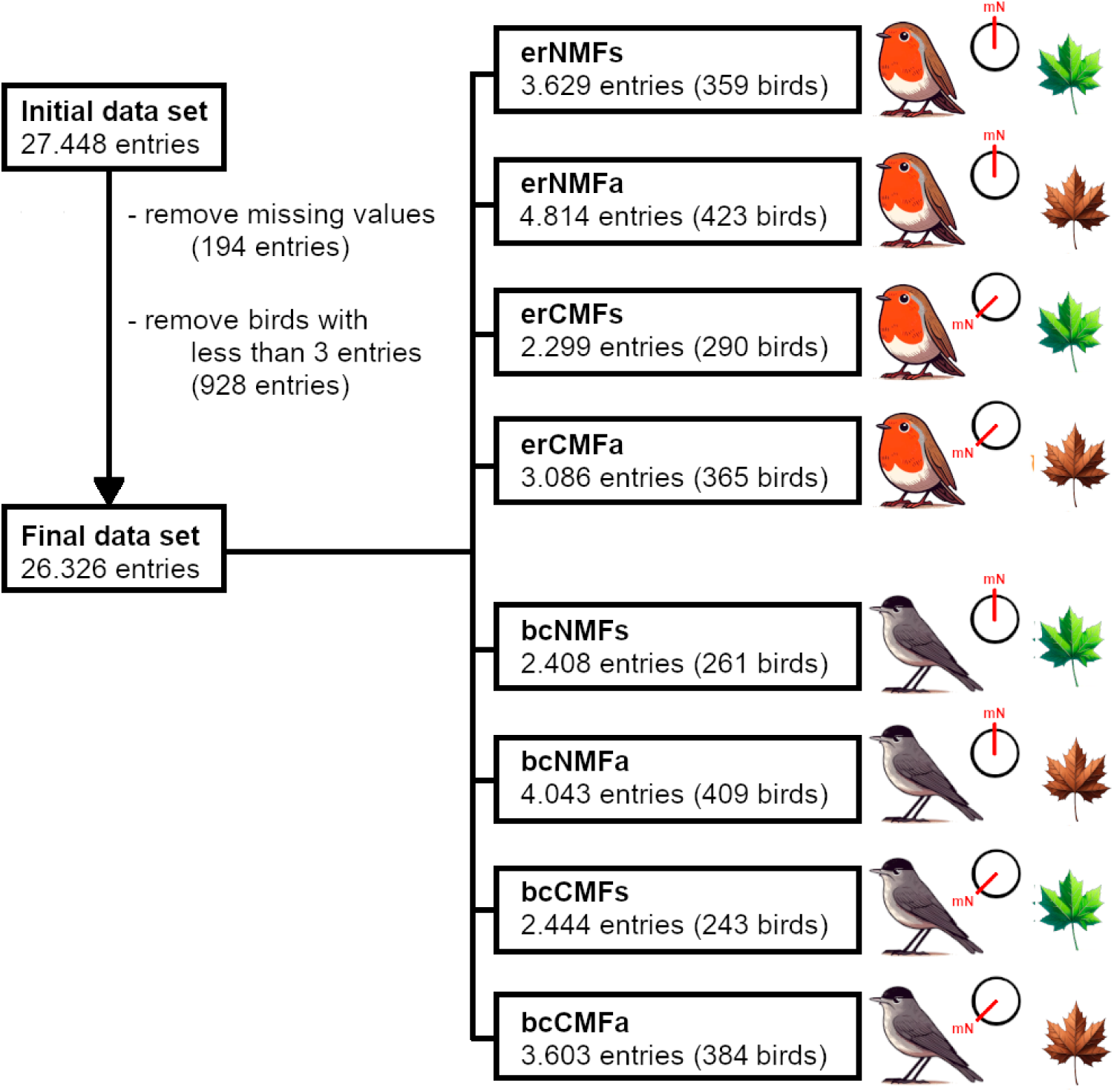
Flow diagram for the selection and categorization of bird data into eight groups. The names of each group describe its particular permutation: European robin (prefix “er-”) vs. Eurasian blackcap (prefix “bc-”); normal magnetic field (NMF) vs. changed magnetic field (CMF); spring (suffix “-s”) vs. autumn (suffix “-a”) migration season. For example, erNMFs corresponds to entries for European robins in NMF during spring, whereas bcCMFa corresponds to entries for Eurasian blackcaps in CMF during autumn.

Among the bird entries, there were several missing values for the activity and the concentration of scratches (194 entries, 0.7% of the total number of entries). These entries were completely excluded from our analysis to ensure consistency. The original experiments have been conducted during the bird’s biannual migratory seasons in spring and autumn, so we pooled entries for each migratory season as February-June (spring) and August-December (autumn), respectively. The compiled data set of Emlen funnel scratched papers included entries for birds that participated in pre-screening (experiments to evaluate the migration motivation of individuals) and actual tests (of pre-screened, motivated birds).

In the beginning of each of the migratory season, pre-screening tests take place to determine (*i*) which birds are motivated to perform orientation behaviour in the funnels and (*ii*) which birds use their magnetic sense to orient inside the funnels. Various parameters can affect the behaviour of birds and ultimately the experimental results. For example, there might be individuals inside the bird groups that are non-migratory, ^61^ or due to time restrictions, some tests may have been conducted outside of the time frame when birds experience migration restlessness.^62,63^ Based on the pre-screening data, the birds classified as active and oriented are selected for the actual experiments. The selection of active and oriented birds is usually based on the consistency of activity and the directedness metric that is recorded for each bird. In general, pre-screening tests are necessary to ensure that only motivated birds participate in the actual tests.

The data set that we compiled by pooling data across all experiments was blind (did not include information regarding) to which birds participated in the actual experiments and which entries correspond to motivated or unmotivated birds. In order to reduce any possible selection bias, our meta-analysis was based on the individual bird directedness (*r_individual_*) and the number of tests (N*_tests_*) regardless of the individual’s activity or direction, Figure 1B. In theory, motivated birds would have a tendency towards a specific direction, and this tendency should be reflected on their individual directedness. To ensure that we had representative statistical data for each individual, only birds with 3 or more entries were considered for the analysis (as was custom in recent studies^50,51^). The final number of entries that was used in our analysis was 26.326 entries (95.9% of the total number of entries), Figure 2.

For our meta-analysis design, the data set was categorized into 8 groups based on bird species (European robin and Eurasian blackcap), magnetic field condition (NMF and CMF) and season (spring and autumn), Figure 2. This classification enabled direct comparison of mean orientation between the eight groups of birds as a function of species, condition, and season. In this analysis, we ignored the minute changes of the earth’s magnetic field over time at Oldenburg, as we assumed that any potential influence cannot be observed in birds’ behaviour in Emlen funnels at present due to the extensive noise in the data. Accordingly, entries from different years have been grouped together.

### Analysis

The Rayleigh test is the most common statistical method to test hypotheses for (normally distributed) circular data and evaluate deviations from uniformity.^64,65^ There are two main variables that can be obtained from a Rayleigh test. First, the *p*-value (*P*) which is the probability criterion for accepting or rejecting the null hypothesis (H*_o_*) that a sample comes from a uniform distribution. Second, the *r* -value (*r*) which is the normalised sum of the mean vector for a set of variables. The *r* -value ranges from 0 to 1, where 0 indicates a random distribution of the entries and 1 suggests that all entries fall on the same spot. Among other parameters, *r* is often used as a directedness criterion (*r_individual_*) for determining which birds are consistently motivated and able to perceive the earth’s magnetic field. ^45,47,49–51,57–60,66^ A typical *r_individual_* cut-off is set at 0.1^45,47,57–60^ or 0.2.^49–51,66^ In these cases, birds that demonstrated *r_individual_* larger than these cut-off values and provided more than a specified number of oriented tests (N*_tests_*) were used for the published statistics. This measure is usually taken to avoid that poorly performing birds (e.g. with few oriented tests or with random orientation *r_individual_ <* 0.2 or 0.1) affect the overall group orientation by contributing a more or less random value. Apart from the *r_individual_* rating, the *P* value can be also used as a selection criterion for individual birds. ^67–69^ Generally, the Rayleigh test captures deviations from uniformity reasonably well, especially when the data is concentrated in one direction. ^65^ Hence, we used this metric to evaluate (*i*) the directedness of each bird (*r_individual_*) and (*ii*) the overall directedness (*r*) for every species, condition and season group in the data set. It is relevant to mention that even though the Rayleigh test is commonly used to analyse orientation data from Emlen funnel experiments, the underlying assumptions for using the test are not always met. For example, a key issue is that the scratches made by each individual bird cannot be assumed to be independent.^52^ For this reason, *r_individual_* is more appropriate for analysing the bird scratches (than *P*), because calculating the sum of the mean orientation vector does not rely on the assumption that the data are independent.

Power analysis is a statistical method used to determine the statistical power of a hypothesis test, which is the probability of correctly rejecting the null hypothesis when it is false. Power analysis is often overlooked during experimental design due to the practical limitations of conducting the experiments, but it should not be ignored when interpreting the statistical analysis outcome. To address this, we performed a power analysis to estimate the minimum sample size (N) that is required to detect an actual effect when performing hypothesis testing using Emlen funnel data. Using appropriate sample sizes reduces the risk of false negative predictions (*β*, Type II error), and increases the statistical confidence in our results. For performing a power analysis, the desired confidence level (*α*) and statistical power need to be defined. Traditionally, *α* is set to 0.05, which is the chance of detecting an effect if there is no such effect (Type I error), and the statistical power (1 - *β*) is set at 0.8, which means that the null hypothesis is correctly rejected in 80% of the cases. The reasons and limitations of this tradition is explained in the statistical literature^70–72^

## Results

### Statistical analysis

First, we assessed the performance of the Rayleigh test for different known distributions to determine the optimal sample size for orientation analysis. The von Mises distribution, see Eq. (1), is used in circular statistics as a correspondence to the normal distribution in linear statistics, making it the most commonly used distribution model for circular data. We used different values for the spread (*k* -value) of the von Mises distribution to generate different distributions and perform controlled sample analyses. Smaller *k* values indicate broader distributions. This simulation approach allows us to thoroughly assess the relationship between the data sampling strategy and statistical analysis outcome, Figure 3A.

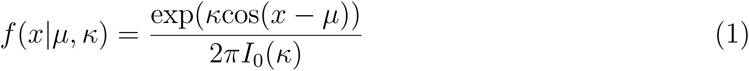

where *µ* is the position of the mean and *I* _0_(*κ*) is a modified Bessel function (of the first kind of order 0).

To explore the performance of the Rayleigh test, we generated data sets by sampling values (N) from different von Mises distributions (by using different *k*), and performed the Rayleigh test for each case. In other words, we have tested the directedness of this circular distribution as a function on its spread (*k*) for different sample sizes (N) to see if they would pass the traditional significance level of *α*=0.05. This controlled simulation shows that even for small sample sizes (N=5 or N=10), highly concentrated distributions (*k>*3) usually give *P* values smaller than the traditional significance level (*α*=0.05), Figure 3B. In contrast, very broad distributions (*k<*1) do not reach the traditional significance level with such small sample sizes. For N=20, most theoretical distributions present a realistic capability of falling below the traditional significance level (*α*=0.05).

For small sample sizes, we find that the Rayleigh concentration parameter (*r*) tends to be elevated, Figure 3C. This trend is even more evident in broader von Mises distributions (*k<*2), and it is present in all sampled distributions. Furthermore, the variation is larger for smaller sample sizes, in particular for broader von Mises distributions (*k<*2). The effects of sample size on *r* has already been reported by Batschelet *et al.*^64^ The trends are displayed in Figure 3C, which shows that distributions for all concentration parameters *k* eventually (and logically) converge to a *r* -value asymptote, given a large enough sample size. Notably, for concentrated distributions (*k>*2), the proximity to this asymptote is reached at considerably lower sample sizes than for less concentrated distributions (*k>*2). The focal aspect of real orientation experiments usually focuses on the deviation of the group orientation from a random distribution (*k* =0). If these theoretical trends are taken as an indication for real experiments, group sample sizes around N = 20 should allow for a differentiation between a random distribution (*k* =0) and broad distributions (*k* =1).

**Figure 3:**
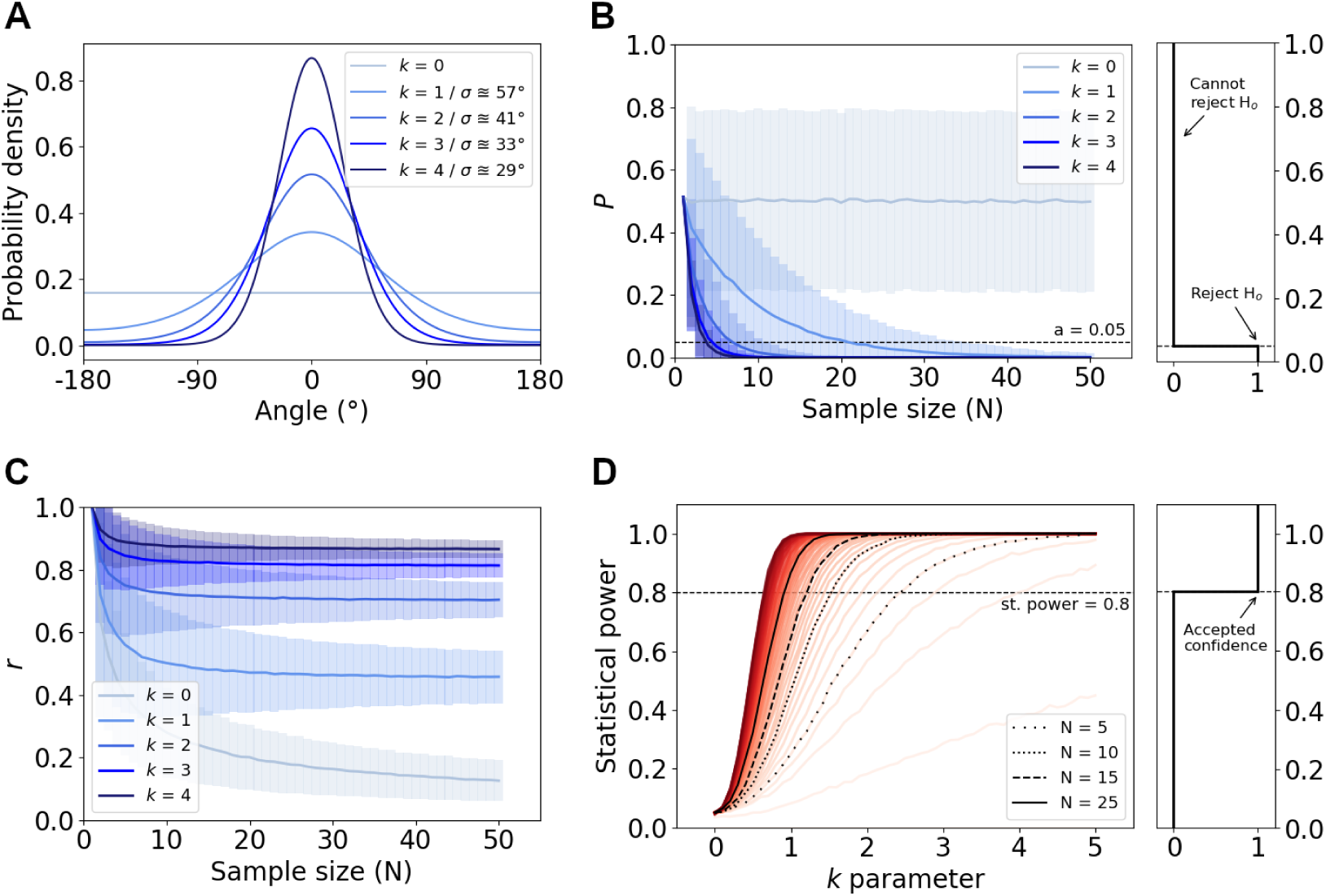
Results from performing the Rayleigh test on different von Mises distributions. A: Von Mises distributions with different spread (*k*), see Eq. (1). Each line corresponds to a different von Mises distribution. The legend includes *k* values with the respective standard deviations (*σ*) from normal distributions. B: *P* -values from the Rayleigh test as a function of sample size (N) for different von Mises distribution spread (*k*). The lines indicate the mean value and the transparent bars represent standard deviation after sampling the von Mises distribution 10.000 times for each combination of *k* and N. The black dashed line highlights the traditional significance level (*α*=0.05). C: *r* values from the Rayleigh test as a function of the sample size for different von Mises distributions. D: Power analysis for rejecting the null hypothesis of uniformity for different von Mises distributions. Estimates are based on 10.000 samples for each combination of *k* and N. The significance level *α* was set to the traditional level of 0.05. The black horizontal dashed line highlights the associated statistical power threshold of 0.8.

The power analysis confirms the aforementioned effects, Figure 3D. Similarly, it also shows that sample sizes around 5 do not provide enough statistical power, when the spread of the data starts getting large (*k<*2). Sample sizes around 15 significantly improve the statistical power, but the results show that ideally larger sample sizes should be used to obtain enough statistical power.

It is important to emphasize that the statistical analysis was performed on data drawn from von Mises distributions, thus the ideal sample sizes apply only for these data. The ideal sample sizes for the von Mises distributions should be used more as a reference, rather than as strict guidelines. Actual experiments tend to produce much noisier data, and larger sample sizes are more likely to capture the real trends, but they can be hard to obtain in reasonable time frames. Furthermore, increasing the sample sizes and the number of tests can raise logistical issues. Regardless, 3R ethical considerations ask researchers to design experiments with sufficient power to be conclusive when the data for calculating power is available. The meta-analysis that we present here enables this.

### Emlen funnels data analysis

The combined data set included in total 26.326 Emlen funnel scratched paper entries from 2.734 individual birds. The first goal was to obtain an estimate of values for the *r* parameter and compare them to ideal von Mises distributions.

In order to analyse the data, we categorised the birds as a function of the number of tests (N*_tests_*; the number of Emlen funnel scratched papers) that were performed for each bird and the individual directedness (*r_individual_*) of each bird, Figure 4. These two parameters, the number of tests and the directedness of each bird, were used to apply the cut-off grouping. Figure 4 explains the cut-off process in the group of European robins tested in normal magnetic field conditions in spring (erNMFs). Each point on these plots represents a group of birds that includes birds with higher values than the N*_tests_* and *r_individual_* cut-offs (red regions in Figure 4A). By increasing the cut-off step-by-step, groups with a higher directedness and a greater number of tests are effectively pooled together. By reducing the cut-off, the bird group sizes increase as birds with lower number of tests and directionality are included in the analysis. Once the groups have been determined, the Rayleigh test is applied to each group, and the directedness (*r*) and *P* value are calculated for each point on the plot, Figure 4B. Then, groups of birds with the same sample sizes (black contour lines) that presented lower *P* values than the significance level (*α <* 0.05) were used for calculating the mean directedness (*r*) for each sample size, Figure 4C. The range of values for N*_tests_* was 3 to 30, and for *r_individual_*, it was 0 to 1, respectively.

**Figure 4:**
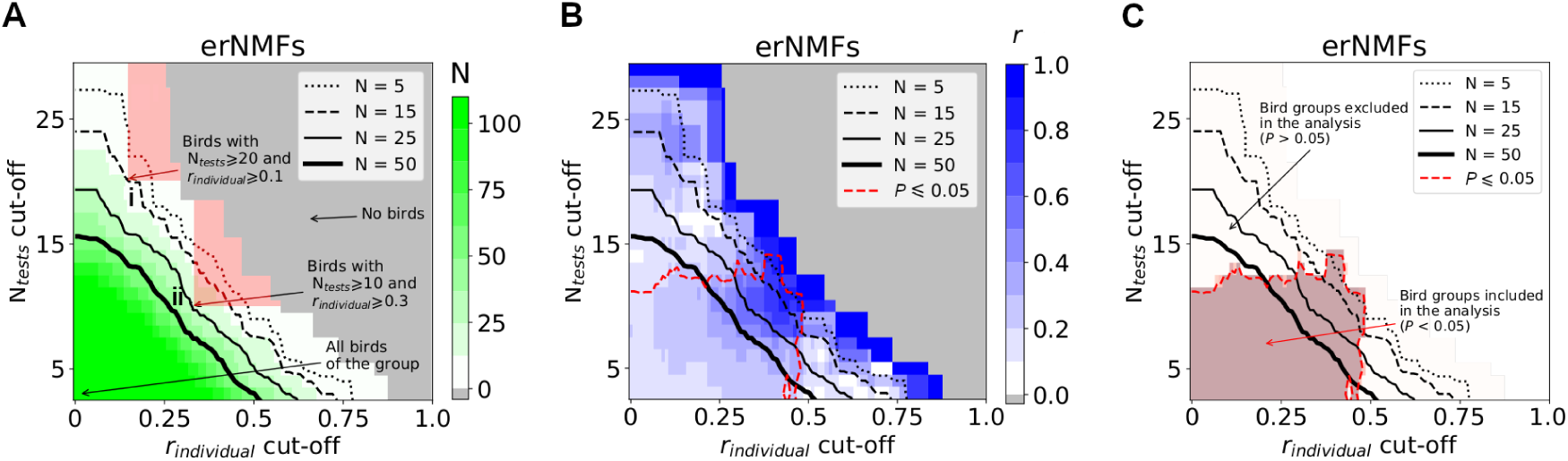
Description of the Emlen data analysis for European robins in the normal magnetic field during spring (erNMFs) group. A: Number of birds as a function of directedness (*r_individual_*) and the number of Emlen funnel tests (N*_tests_*; performd on a signle individual) after applying the cut-off grouping process. The black contour lines indicate the size of each group or, in other words, the number of birds (N = 5, 15, 25 and 50) that met the cut-off criteria in each point of the plot. The red regions highlight the features of birds included in points i and ii. Specifically, i and ii include 15 and 25 birds, respectively. B: Results from the Rayleigh test, *r* as a function of *r_individual_*and N*_tests_* and *P* boundaries. The red contour line highlights the cases for which the *P* of each Rayleigh test is lower than 0.05. C: Bird groups that showed statistically significant deviations from uniformity (*P <* 0.05). The groups within the red region were used for calculating the mean directedness (*r*).

Four main conclusions could be formulated based on the meta-analysis of the existing

Emlen funnel data:

1. The data in the meta-analysis has a high level of noise, as would be expected, given the noisy origin of the data for these behavioural experiments. Groups with a few birds (N*<*5) have high group concentration parameter (*r*), but *r* decreases fast as the group sizes increase. The lower left corner of the plot in Figure 4A, which includes most birds in each group, shows generally small *r* values. The main reason for the low *r* values is the inclusion of pre-screening birds that never reached the actual experiments.
2. Large sample sizes (N*>*50; see the bold contour line in Figure 4B and 4C) present statistically significant deviations from uniformity regardless of the noise. It is sensible that as the sample size increases, small deviations from uniformity are captured easier by the Rayleigh test. This effect was also observed in the analysis of the *P* value for different von Mises distributions, Figure 3B.
3. Individual birds with high directedness (*r_individual_*) did not always give high *r* and small *P* values in the following Rayleigh tests. In these cases, a high *r_individual_* can be explained as an artefact from the small number of tests for the individual birds (N=2-3, Figure 3C), which happened by chance to be consistent in their preferred direction.
4. There are statistically significant deviations from uniformity all along the spectrum of N*_tests_* and *r_individual_*cut-offs. While the example of erNMFs presented a well-defined region with significant deviations from uniformity (area within the red dashed contour lines where *r_individual_>*0.25 and N*_tests_>*5, see Figure 4B and 4C), other analysed groups displayed more spread statistical significant regions and patchier ranges in the cut-off parameters. Plots for all the eight analysis groups are included in the Supporting Information (SI), Figure S1.

The next analysis involves a comparison of the groups that presented statistical significant deviations from uniformity based on the Rayleigh tests and had the same size (number of birds). Ideally, the sample sizes should be the same when comparing different Rayleigh tests, so that all observations have the same weight.^64^ This is not always possible due to practical and logistics issues during the conduct of the behavioural experiments, but the present large (meta-analysis) data set allowed such calculation for different sample sizes. Accordingly, we have calculated the group concentration parameter / directedness (*r*) and the mean orientation of the directed birds for each group for different consistent sample sizes, see Figure 5 and 6, respectively. The same analysis was performed with different statistical significance levels (*α* = 0.1, 0.05 and 0.01) for determining deviations from uniformity, in an effort to assess the robustness of the analysis. The overall trends are maintained regardless of the significance levels, see Figure S2.

**Figure 5:**
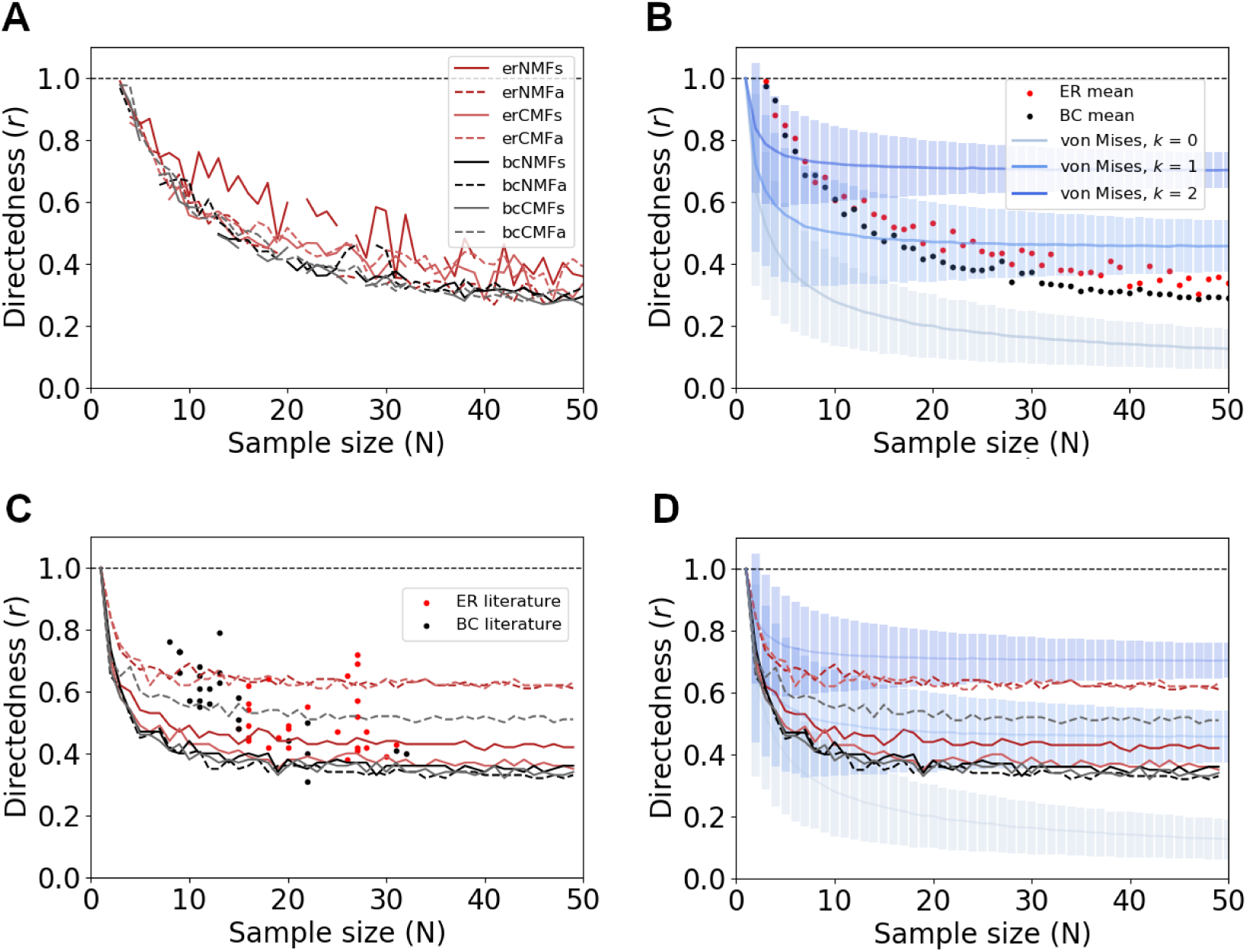
Concentration parameter/directedness (*r*) calculated by performing the Rayleigh test on the different bird groups as a function of the number of birds (N). A: Mean *r* value for the 8 bird groups based on the cut-off grouping scheme and the *P* value from the Rayleigh tests. Each solid red (European robin) and black (Eurasian blackcap) line represents a different species or magnetic field permutation (NMF or CMF) during spring experiments, while the respective dashed lines correspond to autumn experiments. B: Mean *r* value for each bird species and comparison with von Mises distributions. C: Mean *r* value for motivated birds (birds that have passed the pre-selection tests). The *r* value was calculated by sampling randomly bird orientations within each group. The mean *r* was measured by repeating the sampling process 100 times for different N. The dots represent the reported *r* from the respective Emlen funnel experiments included in our sample. D: Comparison between the mean *r* values with von Mises distributions.

**Figure 6:**
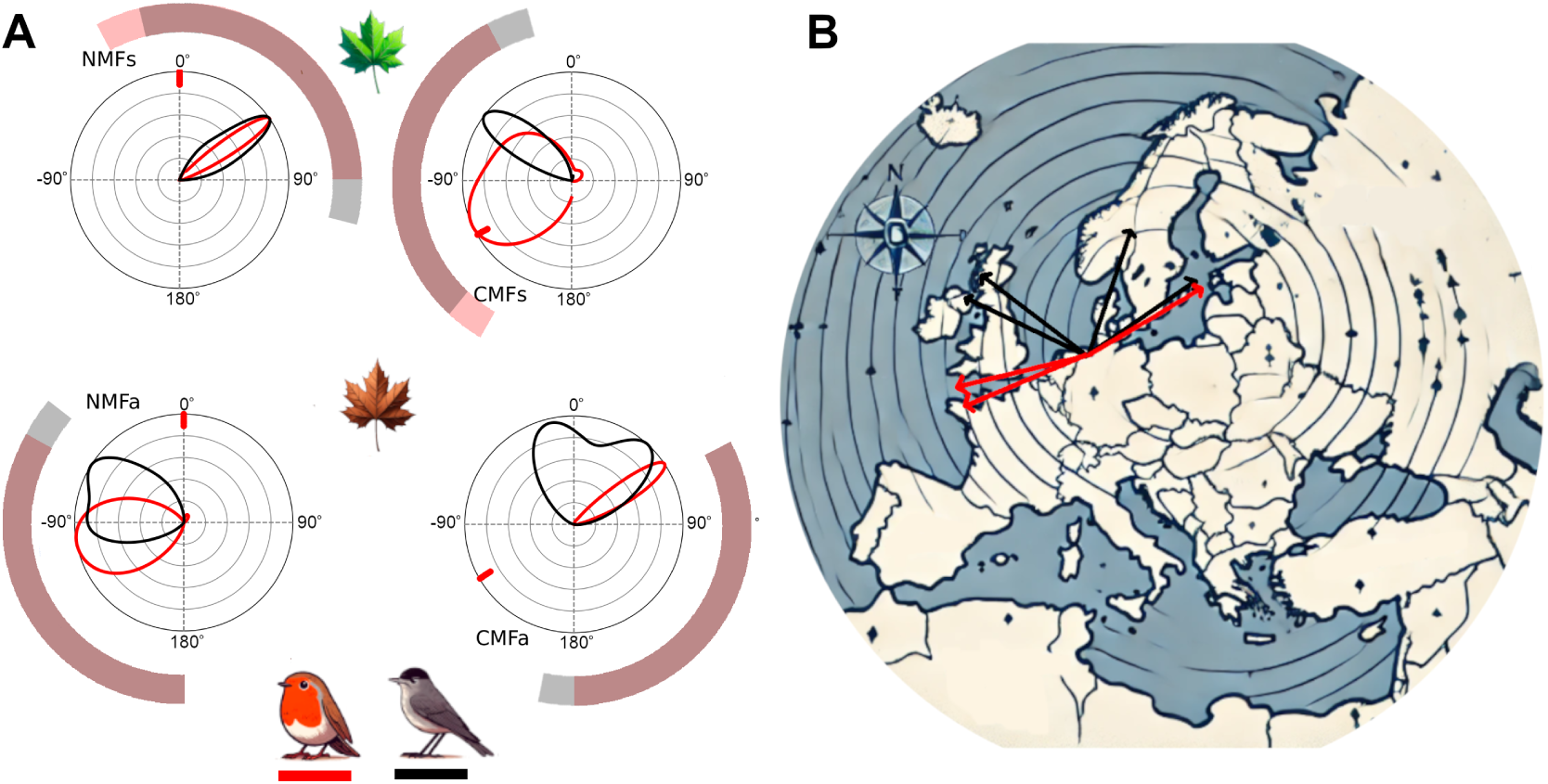
Mean bird orientations for each group considered in this study. A: Density distribution of the mean bird orientation. The transparent red and black slices indicate the expected mean orientation of the European robins and Eurasian blackcaps based on known migration behavior in the wild. This is shown for each season and magnetic field condition. The direction of the magnetic North in each case is highlighted with a red symbol on the outer circle. The density has been normalised for each bird group and condition, separately. B: Superposition of the bird preferred directions on the map to relate their capture and testing location in Oldenburg to their global migration habitat. The preferred direction of European robins and Eurasian blackcaps are shown in red and black, respectively.

The group directedness (*r*) follows a decaying trend similar to the von Mises distributions, but the shape is markedly different, see Figure 5B. Sample sizes (N) around 5 give high *r* values (*∼*0.8-0.9), however *r* is almost half when the sample size is increased to 15 or Since the data were pooled from published studies, the *r* values of the motivated birds (birds that have passed the pre-selection tests) can be calculated compared with the *r* values from the cut-off grouping process, see Figure 5C and 5D. All *r* value lines obtained from motivated birds are more similar to the von Mises distributions, comparison shown in Figure 5D. However, the asymptotes of these lines are quite different among different bird groups, and they range from 0.35 to 0.7. Furthermore, the reported *r* values do not seem to follow the same trends, especially for Eurasian blackcaps, see Figure 5C. These differences could be attributed to selection based on a minimum required *r_individual_* during the pre-tests or the small size of the pooled data that affects the mean *r* value calculation. It is clear from both analyses that the asymptotes at large sample sizes are close to the von Mises distributions with a large spread of data (*k* = 1), which is in line with the intrinsic noise of Emlen funnel data.

As for the mean orientation, there are groups of birds whose mean orientation is roughly the same between species and follows the seasonally appropriate direction (e.g. erNMFs or bcNMFs), but there are also groups that present large spread and deviations (e.g. erCMFs or bcCMFa), see Figure 6. The analysis of the mean bird orientations shows that in most cases the bird groups on average followed the seasonally appropriate orientation directions, see Figure 6B. However, the selection of directed birds based on solely the N*_tests_* and *r_individual_* can lead to artefacts. For example, the presence of birds in the bcCMFa group that were tested many times (large N*_tests_* value), but eventually presented low directionality (small *r_individual_* value) tend to dominate in the tests of statistical significance, see Figure S1. This effect is more evident when the mean orientation is plotted as a function of the N*_tests_* cut-off, Figure S3. Plots for the mean orientation of the oriented bird groups are also included in the Supporting Information (SI), Figure S4.

In order to mathematically model how *r* varies as a function of the bird group sample size (N), we performed a curve fitting analysis, see Figure 7. An exponential function was chosen to represent the mean of the *r* profiles, and the least square method was used to obtain the best fitting curves for the *r* profiles. The fitted curves were compared to the published *r* values from the respective publications to evaluate the performance of our approach for determining directionality, see Figure 7. The superimposition of the modelled *r* values with the published data suggests that our approach captured the main trends well. The *r* profiles for European robins and Eurasian blackcaps are compared with the respective published *r* values of the oriented birds from the studies that we included in our meta-analysis. ^45,47,49–51,57–60^ Two similar functions were obtained for the *r* profiles of the European robins and Eurasian blackcaps, Eq. (2) and (3), respectively.

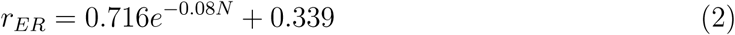

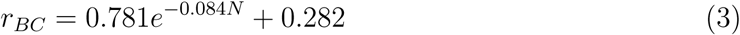

**Figure 7:**
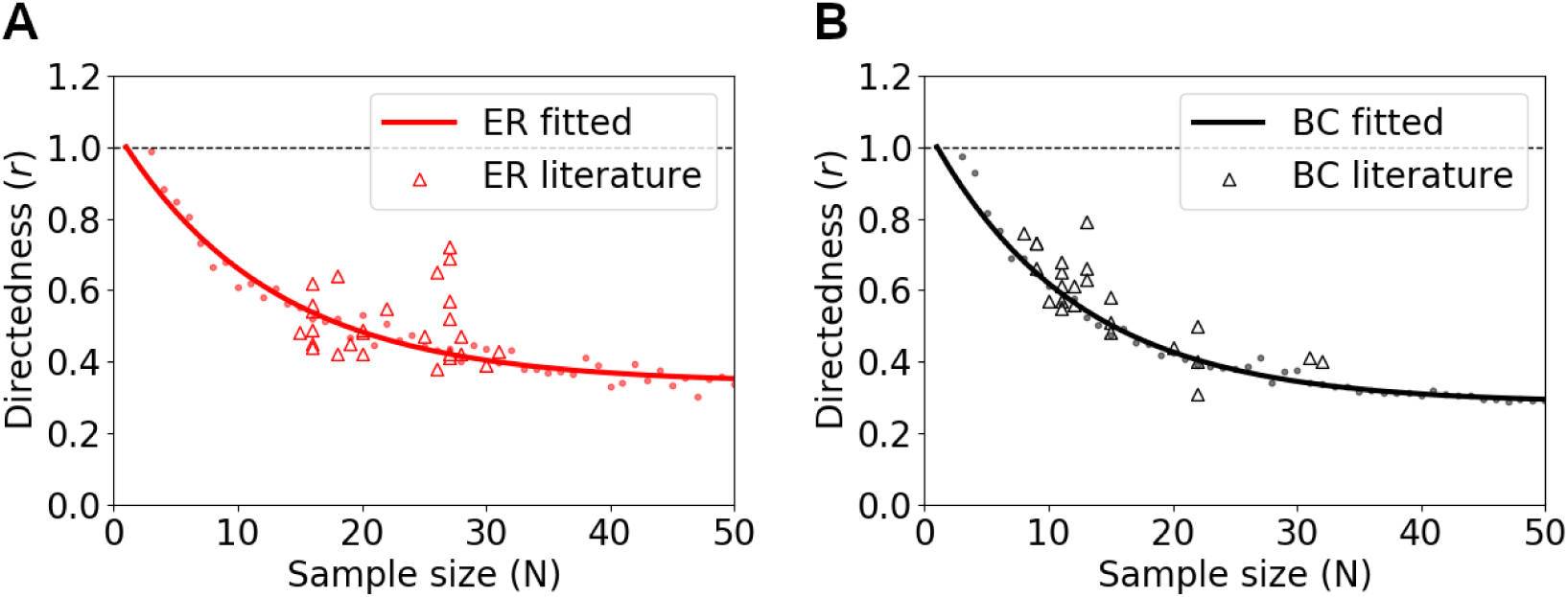
Fitted mean *r* profiles as a function of the sample size and comparison with the published *r* values (triangles) from Emlen funnel experiments from the studies covered by our meta-analysis, European robins (A) and Eurasian blackcaps (B). The mean *r* values (dots) for each bird species has been fitted to an exponential function, Eq. (2) and (3).

The mean absolute errors for the fitting were 0.022 and 0.015 for the European robins and the Eurasian blackcaps, respectively.

The constructed data set has also been analysed without using N*_tests_* and *r_individual_* as cut-off values. Instead, the birds were grouped based on the N*_tests_* and *r_individual_* values. The concentration parameter (*r*), statistical significant deviation from uniformity (*P*), sample size (N) and mean orientation of these groups are included in the Supporting Information (SI), Figure S5. Again, we conclude that the group concentration parameter *r* is larger for the small sample sizes (N*<*5), similar to observations by Batschelet *et al.*^64^ The probability criterion *P* is below the statistical significance level when *r_individual_>*0.2 and 5*<*N*_tests_<*15. In these regions, the mean orientation is usually close to the expected seasonally appropriate direction. Last, there are several outliers in mean orientation, which are mostly groups with small number of tests or small sample sizes.

## Discussion

In this work, we have combined and analysed a large data set of behavioural experiments conducted with European robins and Eurasian blackcaps jumping in Emlen funnels under different magnetic conditions during spring and autumn migration. The jumping behavior was recorded on sensitive paper that the birds scratched with their feet. This meta-analysis enabled us to calculate the average directionality range with respect to the magnetic field for each bird species during spring and autumn migration. Given that magnetic orientation experiments are laborious and challenging to design statistically, due to the high noise in the orientation data, we estimated the minimal sampling requirements for correctly interpreting bird orientation behavior in Emlen funnels using the Raleigh circular statistical analysis adopted by the field.

Our analysis highlights the relationship between the concentration parameter (*r*) and the number of birds tested, the sample size (N). Testing a small number of birds is prone to give extremely large *r* values. Mathematically, we expect a one over sample-size (1/N) like trend boosting *r* values at small samples, which could be negated with appropriate normalization of the concentration parameter in future work.

A successful strategy for reducing noise in orientation data is to remove non-migration motivated birds that show little activity or are not oriented in a consistent direction. In the directed bird groups selected via such pretests, we find that *r* shows the same trend in both species and demonstrates that the birds are indeed oriented. However, in both species this trend also deviates from how *r* varies according to the widely adopted von Mises distributions, showing the underlying assumptions of the Rayleigh test are not strictly met in bird magnetic orientation behavior. The trends in both bird groups are similar to the published values, which helps confirm that our approach captured the differences between directed and non-directed birds as reported before. However, we did note that the mean orientation does not always follow the expected seasonally appropriate migration orientation trend. This raises new research questions about why birds tested in Emlen funnels sometimes orient in a mean direction different from their expected orientation direction in the wild. These more fundamental questions cannot be resolved with a minimal cut-off requirement for the number of tests needed to correctly determine if birds are oriented using the Rayleigh test. Defining better generalizable pretest criteria, ideally grounded in the mathematics of the statistical model used, for discriminating motivated and unmotivated birds could maybe advance the field’s ability to analyze Emlen funnel data.

According to our statistical analyses and meta-analysis, ideal sample sizes for future Emlen funnel studies should range at least between 15–25 to maintain a small probability of statistical error for a species, regardless of the underlying data distribution. These sample sizes are expected to be large enough to provide statistical confidence even when dealing with noisy data, such as behavioural data from Emlen funnels. Small sample sizes are bound to give higher *r* values that are not necessarily close to the asymptote of the actual directedness profiles or the expected seasonally appropriate direction. The nature of the experiments, the inherent noise, the practical limitations and ethical issues do not offer any easy solution to address the sample size effect, but consistency and reproducibility should be of major importance when designing Emlen funnel experiments.

Overall, this study constitutes the largest meta-analysis of laboratory-based data on the magnetic orientation of migratory birds. Our results provide a realistic range of values for the expected directness of birds in Emlen funnels that could be used as a reference for future studies.

## Acknowledgements

The authors would like to thank the Volkswagen Foundation (Lichtenberg Professorship to I.A.S.), the Deutsche Forschungsgemeinschaft (SFB 1372: Magnetoreception and Navigation in Vertebrates, no. 395940726 to I.A.S. and H.M.; TRR386/1-2023 Hyperpolarization in molecular systems HYP*MOL, no 514664767 to I.A.S., and FR 2715/6-1 PreCePT), and the Ministry for Science and Culture of Lower Saxony (Simulations Meet Experiments on the Nanoscale: Opening up the Quantum World to Artificial Intelligence (SMART) and Dynamik auf der Nanoskala: Von kohärenten Elementarprozessen zur Funktionalität (DyNano)). Computational resources for the simulations were provided by the CARL Cluster at the Carl-von-Ossietzky University Oldenburg, which is supported by the DFG and the ministry for science and culture of Lower Saxony. The authors gratefully acknowledge the computing time made available to them on the high-performance computers HLRN-IV at GWDG at the NHR Centers NHR@Göttingen. These Centers are jointly supported by the Federal Ministry of Education and Research and the state governments participating in the NHR.

