## Supplementary Information for "Directionality range in Emlen funnels"

### Supporting Information

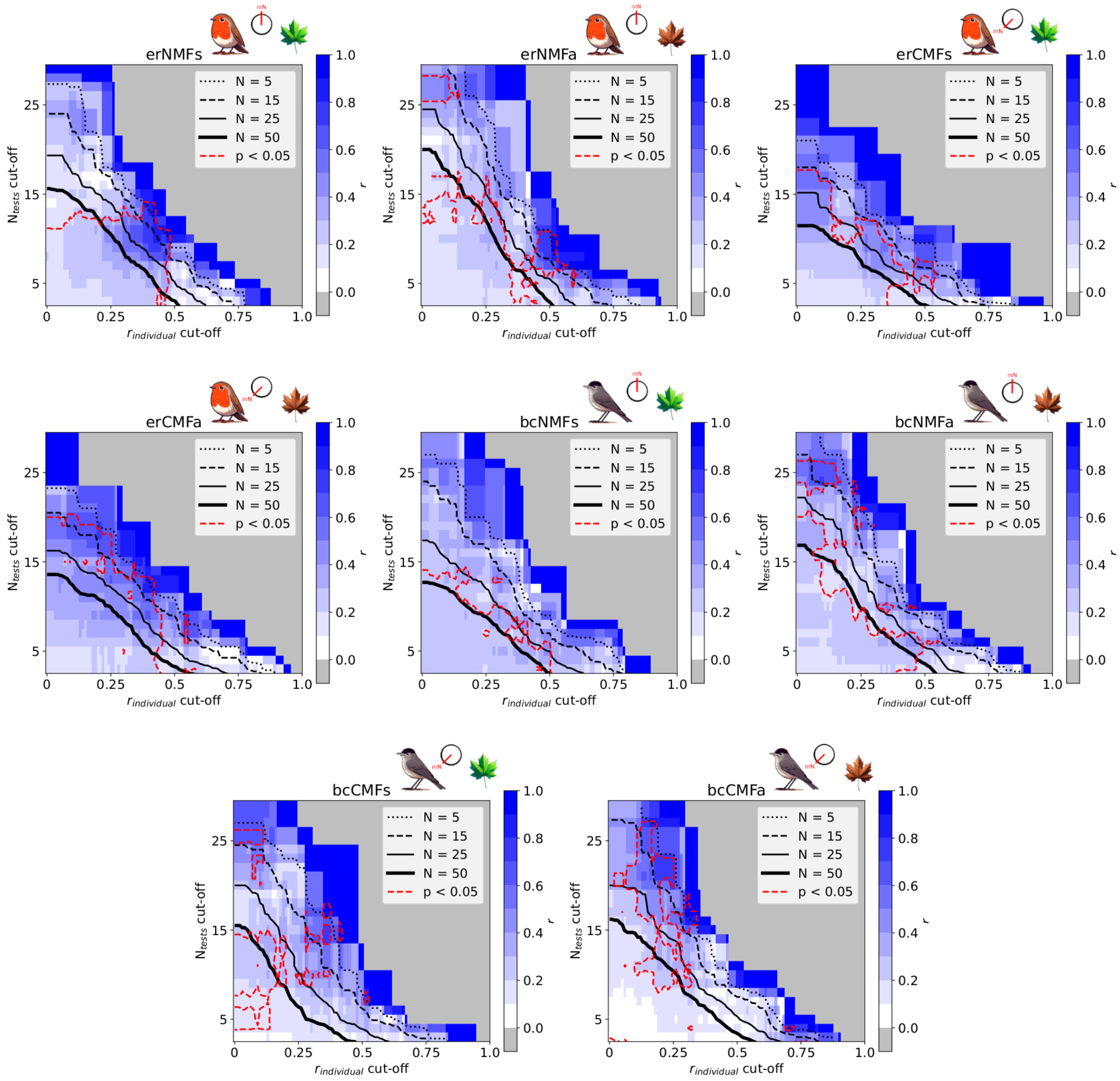


Figure S1: Concentration parameter ($r$) as a function of directedness ($r_{individual}$) and the number of Emlen funnel tests ($N_{tests}$) per bird for each group. The black contour lines indicate the boundaries for different sample sizes (5, 15, 25 and 50) for the $r$ calculation and the red contour lines highlight the cases when the $P$ was lower than 0.05 for each Rayleigh test. The light grey areas indicate the regions with no birds.

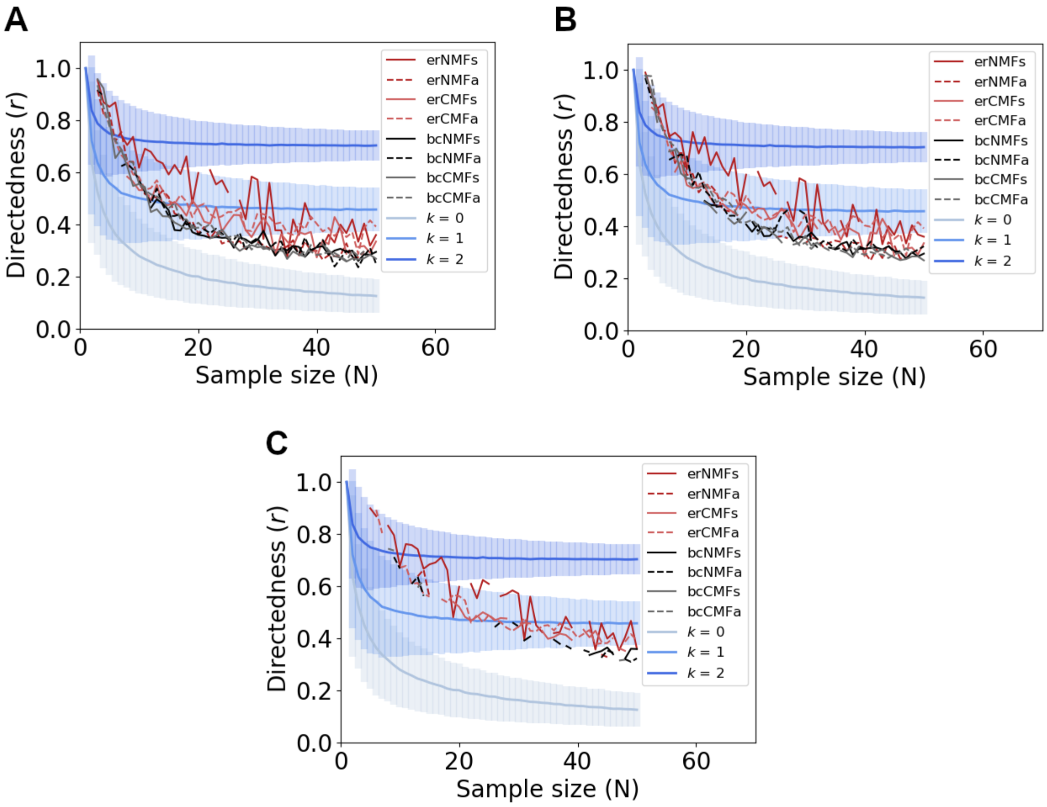


Figure S2: Directedness ($r$) calculated by performing the Rayleigh test on the different bird groups as a function of the number of birds (N). The statistical significance for each plot is different, 0.1 (A), 0.05 (B) and 0.01 (C). The trends are quite similar regardless of the significance level. However, decreasing the significance level to 0.01 (C) results in an small increase of the $r$ values due to the exclusion of the less oriented birds.

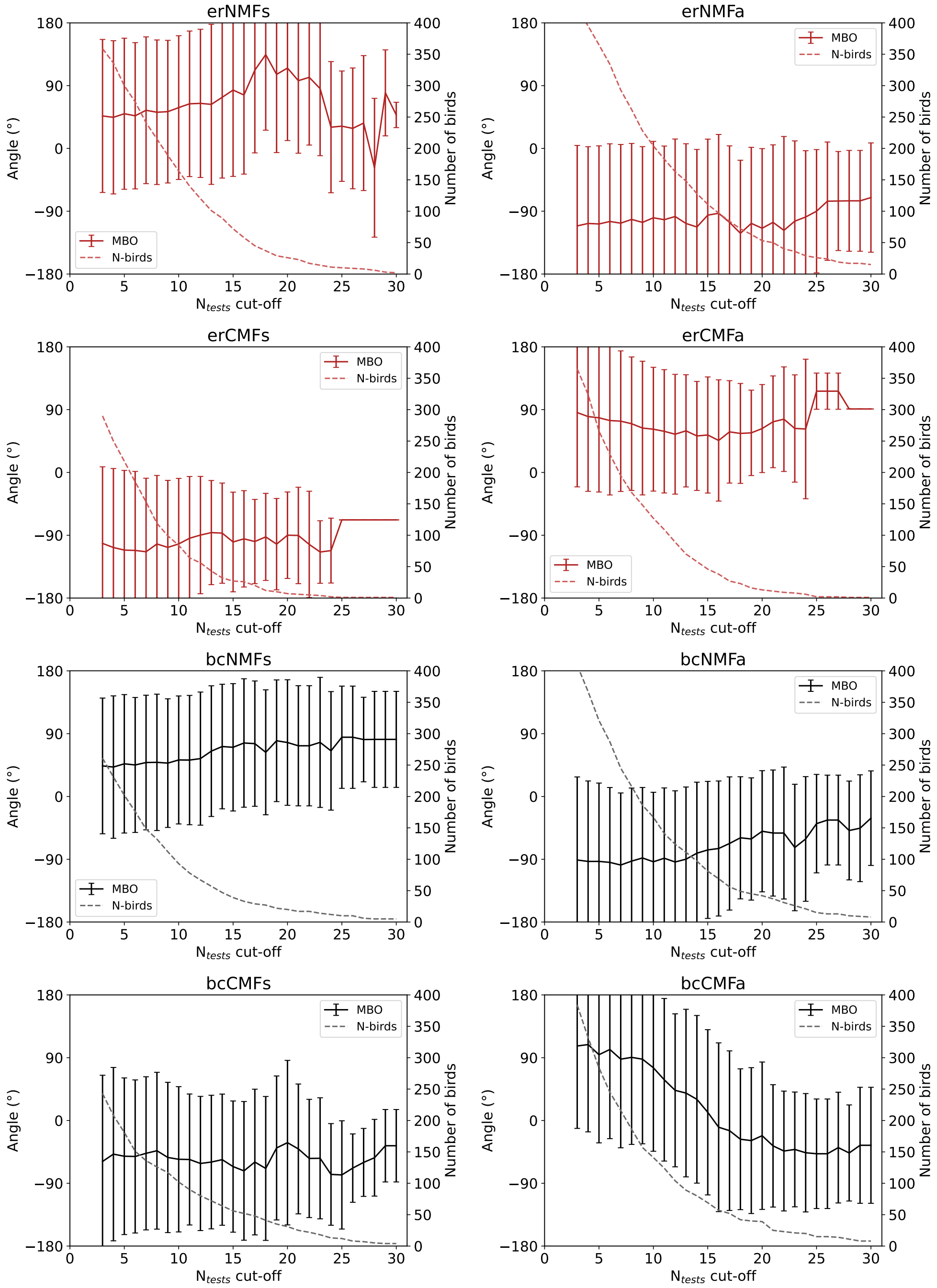


Figure S3: Mean orientation of birds as a function of the $N_{tests}$ cut-off for each group. The error bars correspond to standard deviation. The dashed lines indicate the number of birds for calculating the mean values.

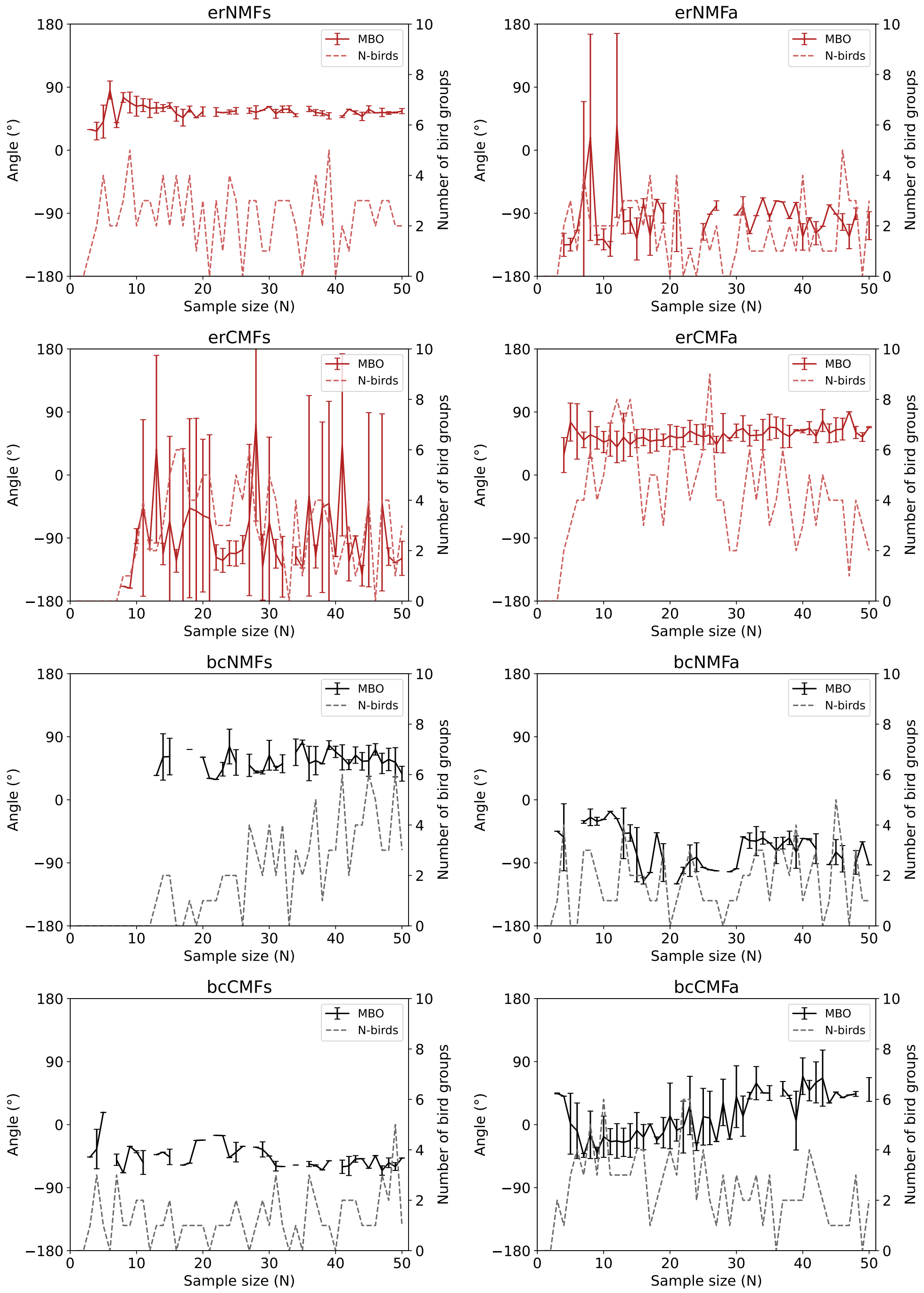


Figure S4: Mean orientation of oriented birds (based on our directedness selection criteria) as a function of the sample size (N) for each group. The error bars correspond to standard deviation. The dashed lines indicate the number of birds for calculating the mean values. Some bird groups present deviations in their mean orientation, but overall the seasonally appropriate orientation is captured especially when sample sizes are large.

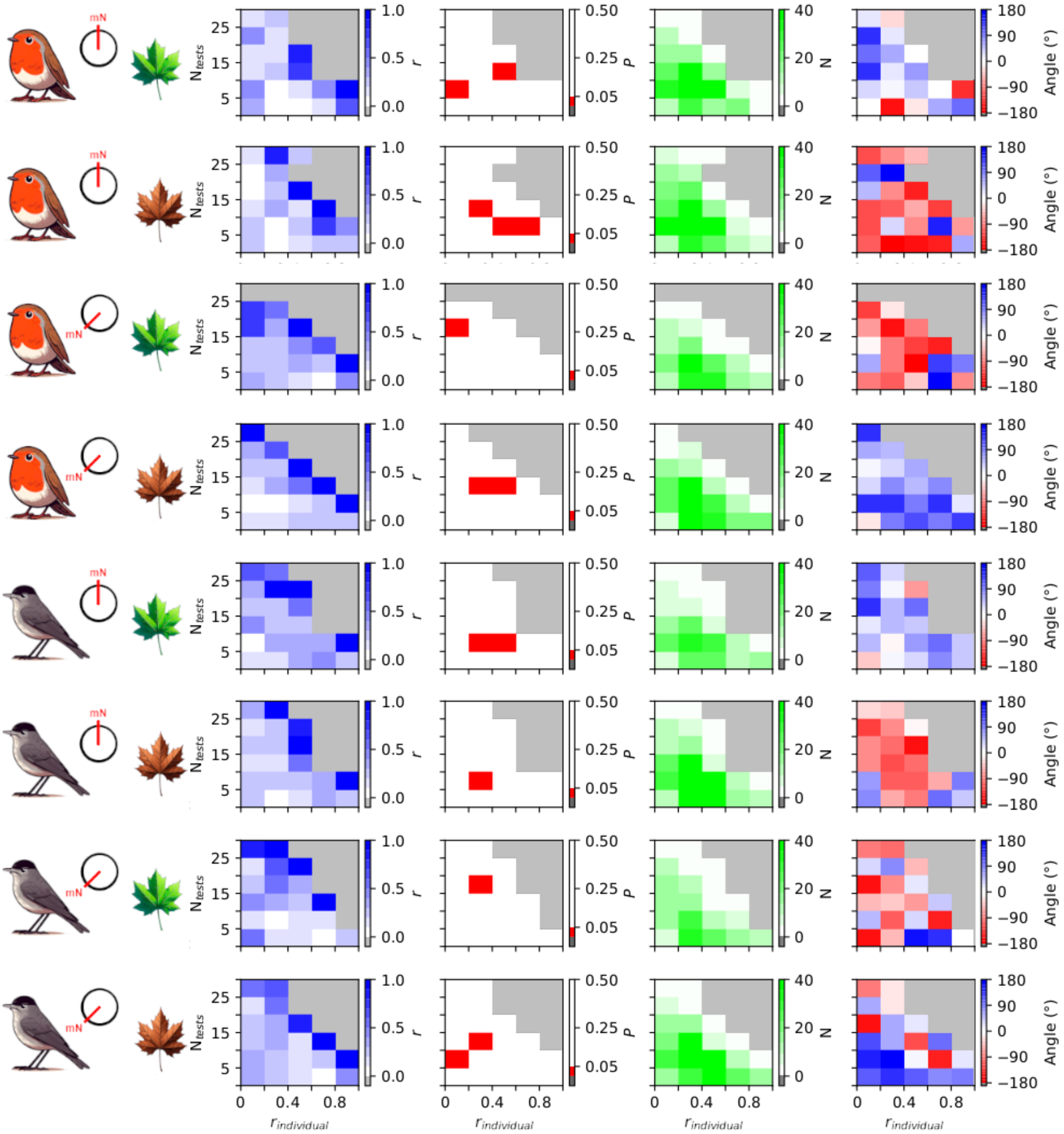


Figure S5: Concentration parameter ($r$), probability criterion ($P$), sample size (N) and mean bird orientation for different bird groups based on $r_{individual}$ and $N_{tests}$ subgroups. The light grey areas indicate the regions with no birds.
